# Drug-induced resistance evolution necessitates less aggressive treatment

**DOI:** 10.1101/2020.10.07.330134

**Authors:** Teemu Kuosmanen, Johannes Cairns, Robert Noble, Niko Beerenwinkel, Tommi Mononen, Ville Mustonen

## Abstract

Evolution of drug resistance to anticancer, antimicrobial and antiviral therapies is widespread among cancer and pathogen cell populations. Classical theory posits strictly that genetic and phenotypic variation is generated in evolving populations independently of the selection pressure. However, recent experimental findings among antimicrobial agents, traditional cytotoxic chemotherapies and targeted cancer therapies suggest that treatment not only imposes selection but also affects the rate of adaptation via altered mutational processes. Here we analyze a model with drug-induced increase in mutation rate and explore its consequences for treatment optimization. We argue that the true biological cost of treatment is not limited to the harmful side-effects, but instead realizes even more profoundly by fundamentally changing the underlying eco-evolutionary dynamics within the microenvironment. Factoring in such costs (or collateral damage) of control is at the core of successful therapy design and can unify different evolution-based approaches to therapy optimization. Using the concept of evolutionary rescue, we formulate the treatment as an optimal control problem and solve the optimal elimination strategy, which minimizes the probability of evolutionary rescue. Our solution exploits a trade-off, where increasing the drug concentration has two opposing effects. On the one hand, it reduces *de novo* mutations by decreasing the size of the target cell population faster; on the other hand, a higher dosage generates a surplus of treatment-induced mutations. We show that aggressive elimination strategies, which aim at eradication as fast as possible and which represent the current standard of care, can be detrimental even with modest drug-induced increases (fold change *≤*10) to the baseline mutation rate. Our findings highlight the importance of dose dependencies in resistance evolution and motivate further investigation of the mutagenicity and other hidden collateral costs of therapies that promote resistance.

**Author summary:** The evolution of drug resistance is a particularly problematic and frequent outcome of cancer and antimicrobial therapies. Recent research suggests that these treatments may enhance the evolvability of the target population not only via inducing intense selection pressures but also via altering the underlying mutational processes. Here we investigate the consequences of such drug-induced evolution by considering a mathematical model with explicitly dose-dependent mutation rate. We identify, characterize and exploit a trade-off between decreasing the target population size as fast as possible and generating a surplus of treatment-induced *de novo* mutations. By formulating the treatment as an optimal control problem over the evolution of the target population, we find the optimal treatment strategy, which minimizes the probability of evolutionary rescue. We show that this probability changes non-monotonically with the cumulative drug concentration and is minimized at an intermediate dosage. Our results are immediately amenable to experimental investigation and motivate further study of the various mutagenic and other hidden collateral costs of treatment. Taken together, our results add to the ongoing criticism of the standard practice of administering aggressive, high-dose therapies and stimulate further clinical trials on alternative treatment strategies.

## Introduction

The formation of cancer and emergence of antimicrobial resistance (AMR) are notorious examples of fast paced evolution. Modern medicine has developed various drugs to target cancer and pathogen cell populations with the aim to drive them to extinction using aggressive, high-dose therapies. However, these treatments frequently fail due to drug resistance, a phenomenon where the drug loses its desired pharmacodynamical effects. The emergence of drug resistance is the consequence of evolution which continues also during the treatment. Indeed, the administration of treatment represents a major switch point in the evolutionary trajectories of these populations, initiating a rapid phase of human-induced evolution.

The most desirable consequence of this human-induced evolution is the decay of the drug-sensitive target population. The key question in such a situation is whether adaptive evolution can happen fast enough to save the population from extinction. If the population is saved, we say that an *evolutionary rescue* has occurred [1]. Introduced first in the field of conservation biology, where the objective has been to design the most efficient intervention programs to save endangered species from going extinct, the concept of evolutionary rescue can be readily applied also to the study of drug resistance [2, 3] with the opposite goal in mind.

Evolutionary rescue can occur either by standing variation or by *de novo* mutation. Rescue by standing variation corresponds to the intrinsic resistance model in which the population is sufficiently diverse to contain individuals that can survive in the changed environment. Rescue by *de novo* mutation, on the other hand, corresponds to the acquired resistance model in which (partially-)resistant individuals are created by mutational processes after the initiation of therapy. The deeply rooted paradigm of administering treatment as aggressively as possible to maximize cell kill [4] has its origin in the somatic mutation theory of drug resistance [5], where it is assumed that rescue mutations arise spontaneously and independently of the treatment. The rationale of such aggressive elimination therapies is then to maximize the probability of cure by eradicating the population as fast as possible thus minimizing the rescue window during which mutations can occur and save the population.

However, the gain of population decay, comes necessarily with a cost, which realizes as collateral damage at various scales. The most obvious examples of such damage are the clinical side-effects of the treatment, which often result from the off-target exposure to the drug. For example, traditional anticancer therapies hit also healthy tissues while antimicrobial agents negatively affect the natural gut microbiome. The detrimental side-effects experienced by the patient yield a toxicity constraint which have led to the maximum tolerated dose (MTD) paradigm [6], where treatment is predominantly administered at the highest cumulative dose possible given the toxicity constraint.

Recent research and experimental evidence suggest that the true biological cost of the treatment is not limited to the harmful side-effects, but instead realize even more profoundly by fundamentally altering the underlying eco-evolutionary dynamics within the microenvironment. The harsh selection pressure induced by the treatment not only leads to the decay of the sensitive cells, but it can also enhance the growth opportunities of the pre-existing or emerged resistant cells, a well-known phenomenon of competitive release [4, 7]. In these cases, aggressive chemotherapy only accelerates the population’s evolution towards treatment resistance as by removing the competing sensitive cells the resistant cells have even more resources to reoccupy the niche leading to relapse.

This problem has then motivated various authors to suggest so-called containment strategies which use the minimal amount of control to keep the population burden in check while deliberately maintaining sensitive cells to competitively suppress the growth of the existing resistant cells as a form of ecological control [8–11]. Competitive release represents an important ‘ecological collateral damage’ of treatment, which promotes the emergence of drug resistance and leads to treatment failure. Besides the altered competition dynamics, beneficial rescue mutations may become enriched in the off-target species and promote the emergence of AMR by means such as horizontal gene-transfer [12].

In addition to the extensive ecological consequences, the treatment may also induce changes to the intrinsic dynamics of the target cells other than Darwinian selection. Such ‘evolutionary collateral damage’ can realize, for example, by the treatment enhancing the evolvability of the population. Classical theory, based on the famous Luria-Delbrück experiments [13] and somatic mutation theory of drug resistance, posits strictly that genetic and phenotypic variation is generated independently of the selection pressure. In contrast, recent experimental evidence suggests that the therapies themselves may affect the way variation is generated. Studies in bacteria demonstrate that stress alone can increase the genome-wide mutation rate as adaptation to a changing environment (adaptive mutability), driven by switch to more error-prone DNA repair mechanisms [14]. Recently, similar findings were reported also in cancer in the context of targeted cytostatic therapies [15, 16]. Higher levels of reactive oxygen species (ROS) during the treatment may serve as another mechanism which increases the mutation rate [17]. These are examples of fundamentally random drug-induced effects, where only the probability of acquiring mutations changes as a function of dose. On the other hand, conventional genotoxic chemotherapies may also cause specific drug-induced mutations, such as the reported distinct mutational signatures of platinum-based therapies [18].

Secondly, the treatment can also increase the *phenotypic* mutation rate directly as certain resistance mechanisms can be activated even without the need for a mutation using epigenetic regulation in a post-adaptive manner. Cancer cells – especially stem-like cancer cells – can exhibit diverse phenotypic plasticity, dynamically responding to changes in their environment. Cells can enhance their survival via life-history trade-offs by reallocating resources normally devoted to proliferation [19]. Such quiescent, drug-tolerant cells can act as a reservoir from which permanently resistant cells can emerge via further genetic mutations, or alternatively, revert back to active proliferation upon treatment discontinuation [20].

Finally, the phenotypic mutation rate can also change due to the dose-dependency of the required mutational targets [21]. Higher levels of stress can decrease the phenotypic mutation rate as the number of mutational targets required for the resistant phenotype may increase. Alternatively, fewer mutational targets may be needed to become selectable as the proportion of beneficial mutations may increase as the selection pressure becomes harsher. This may promote the emergence of drug resistance by gradual step-wise adaptation, especially in drug-sanctuaries [22]. All these findings provide compelling evidence that the genotypic/phenotypic mutation rate, and hence the rate of adaptation to the stressful environment, is strongly dose-dependent.

The outlined collateral damage occurring at various scales ranging from the whole patient to the microenvironment of the target cells greatly complicate the combat against drug resistance and require the integration of ecological and evolutionary dynamics into therapy design. Eco-evolutionary control has to further factor in the underlying biological mechanism of control [23]. Therapy is often based on biomolecular interactions, such as drug–target or antibody–antigen binding [24]. The biophysics of molecular binding dictate a finite control leverage, for an example, as seen in the Hill-function type pharmacodynamics which cause a saturation of the drug’s effect. However, the different collateral damage caused by the drug do not saturate in general, but can keep increasing with the dosage, or saturate at a different concentration. These observations point to a great scope in designing treatments using eco-evolutionary control theory [23].

Majority of previous treatment optimization models have focused on optimizing the delivery mode with respect to some toxicity constraint (see e.g. [25]). The “second-wave” of treatment optimization has focused on the issue of competitive release and the investigation of various containment strategies [11, 26]. Here we investigate the consequences of evolutionary collateral damage, as realized by drug-induced resistance evolution, on treatment selection using the rigorous methods of optimal control theory [27] (see Fig. 1).

**Fig 1.**
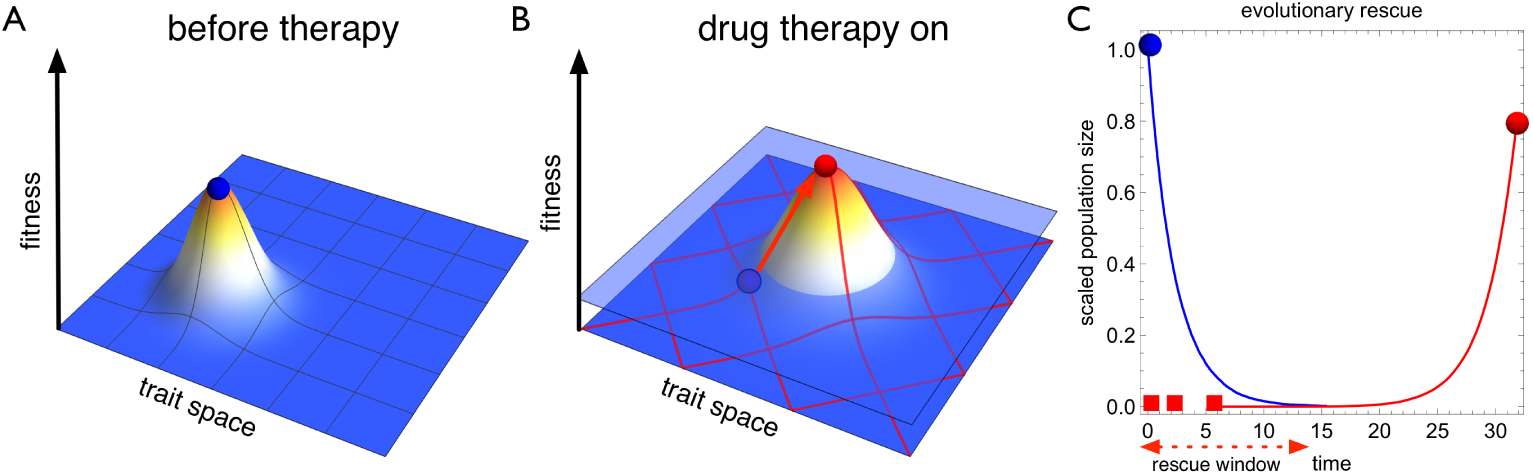
Drug-induced mutations realise an evolutionary collateral cost of therapy. **A** Before therapy the target cell population (blue circle) is well-adapted to its microenvironment. **B** Initiation of control (drug therapy) drastically changes the growth conditions of the target population pushing it below zero level of growth (light blue plane). At the same time opportunities to adapt to the new conditions create a selection pressure for resistance to evolve. Furthermore, the therapy can change the mutational wiring both qualitatively and quantitatively (red mesh). The effect of therapy on the mutational processes represent an evolutionary collateral damage of control, which can expedite the emergence of resistance (red arrow and circle). **C** The treatment eliminates the sensitive cell population (blue) during the rescue window but an evolutionary rescue can occur if a resistant mutant (red squares) manages to successfully establish. Here we derive optimal treatment strategies that minimise the probability of evolutionary rescue while taking into account drug-induced mutations.

Only few previous theoretical studies of drug resistance have explicitly accounted for dose-dependent mutation rates [28, 29]. Liu *et al*. find that the optimal delivery mode of the MTD-strategy is robust against changes in the mutation rate in a model for targeted cytostatic cancer therapy. Greene *et al*. on the other hand show that the dose-dependent mutation rate may have a significant impact on which delivery mode is preferable. In contrast, we solve the optimal elimination strategy, which minimizes the probability of an evolutionary rescue, and demonstrate the potential to improve treatment outcomes by reducing the total cumulative dosage administered.

We identify and exploit a trade-off, where increasing the dosage on the one hand reduces *de novo* mutations by decreasing the target population size faster, but at the expense of simultaneously generating a surplus of treatment-induced mutations. By simulating virtual treatments *in silico*, we show that the probability of cure changes non-monotonically as a function of the drug concentration and is maximized at an intermediate dosage. Our results highlight the importance of dose-dependencies in resistance evolution and help to redefine the precise evolutionary objectives of the treatment, providing a framework for systematic therapy optimisation.

## Results

### Dose-dependent mutation rate introduces a trade-off between mutation intensity and target population decay

Here we set the therapy optimisation problem by formulating the specific control objectives that we want to achieve while factoring in the eco-evolutionary dynamics of the target population. We further demonstrate how drug-induced mutations affect realising these goals.

Figure 1C provides a schematic example of an evolutionary rescue: first, the population rapidly declines as the treatment eradicates sensitive cells. Cells can however acquire mutations that reduce their sensitivity to the drug. The emergent resistant cells are strongly favoured by natural selection but can nevertheless be lost due to stochastic extinction [30]. If resistant cells manage to establish, they will soon repopulate the tumor niche and rescue the population from extinction.

We use the term *rescue window* for the initial treatment period during which sensitive cells can acquire mutations and the population can be rescued. We model the acquisition of rescue mutants by a time-inhomogeneous Poisson process during the rescue window, where the rate of gaining a new mutant at time *t* is given by the product of the (sensitive) cell population size *S*(*t, u*(*t*)) and the phenotypic mutation rate *µ*(*u*(*t*)). Both of these factors depend on the drug concentration *u*(*t*), as we explicitly take into account drug-induced effects.

Because the growth of resistant cells originates from a single cell, we must use stochastic population dynamics to model the growth of small populations that have a considerable extinction risk due to inherent stochastic fluctuations. The stochastic extinction risk for a simple birth-death process founded by a single resistant (subscript *R*) cell is given by 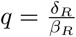 (see e.g. [31]), where *δ*_*R*_ and *β*_*R*_ are the intrinsic death and birth rates respectively, leading to the net growth rate *r*_*R*_ = *β*_*R*_ *− δ*_*R*_. For simplicity, we assume that this stochastic extinction probability is a constant property of the resistant cell and does not depend on the control variable *u* directly (by assuming complete resistance) or indirectly via sensitive cell density (see Materials and Methods). If control can also be exerted to the resistant cells (partial sensitivity), the optimal control can further capitalize on this establishment probability.

Now consider an *elimination strategy u* : [0, *T*] *→ 𝒰* = [0, *u*_max_], which gives the desired drug concentration over the treatment period. Here the end-time *T* is assumed fixed and we allow it to be long enough so that there are no sensitive cells left at the end, *S*(*T*) = 0. With these assumptions, the intensity of the Poisson process, or the total cumulative rate of generating rescue mutations, is given by

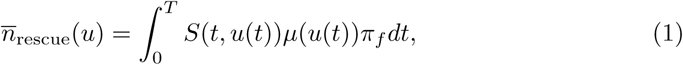

where *π*_*f*_ = 1*− q* is the probability of establishment of a new mutant. This quantity corresponds to the expected number of successfully established rescue mutants generated during the treatment period. The probability of an evolutionary rescue by *de novo* mutation is then [2]

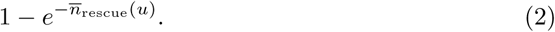

The exponential term is just the zero class of the Poisson distribution and hence the complement of this gives the probability that there is at least one cell which survived the rescue window. A suitable objective of the treatment is then to minimise this quantity, which is equivalent to maximising the extinction probability of the target population. Since the probability of establishment is here just a constant, the objective functional for the optimal control problem reduces to

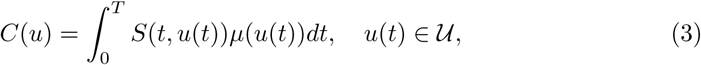

which corresponds to the expected number of mutant establishment attempts. The discussed control problem is then to find the optimal elimination strategy, which minimizes the cost functional above from the space of all (Lebesque integrable) functions over the treatment period.

Because the probability of gaining mutations is proportional to the population size, a characteristic feature of the rescue window is that the probability of an evolutionary rescue sharply decreases with decreasing population size. This phenomenon corresponds to the classical somatic mutation theory of drug resistance and justifies the MTD strategy which aims to minimise the probability of an evolutionary rescue by making the rescue window as short as possible. Indeed, suppose that the phenotypic mutation rate *µ* is independent of the control variable. Then, the MTD solution *u ≡ u*_max_ is trivially the optimal treatment strategy and the treatment can be optimised only with respect to the delivery mode that satisfies the cumulative toxicity constraint. However, if *µ*′ (*u*) *>* 0, then clearly the MTD solution is generally not optimal, because now the population size can be decreased only at the expense of increasing the mutation rate leading to an interesting and potentially exploitable trade-off, where the optimisation can be done also with respect to the cumulative drug concentration.

Figure 2 shows the intensity at which rescue mutations are generated during treatment. If no treatment is administered (*u* = 0), the population grows to its carrying capacity and generates rescue mutants with the constant baseline mutation rate *µ*_0_. When treatment is administered (*u*(*t*) *>* 0), the mutation probability sharply decreases as the population size decreases. However, the drug-induced effects create a mutational peak in the beginning, where the probability of rescue mutations rises above the baseline mutation rate as control is being applied to a large population size. Higher doses lead to a higher early mutational peak, but the probability of rescue mutations decreases faster as the sensitive cell population diminishes faster than at lower doses. Lower doses on the other hand have a lower early mutational peak, but as it takes longer to eliminate the sensitive cells, the mutation probability decreases more slowly, thus prolonging the rescue window. The optimal strategy, which minimises the total cumulative rate of generating rescue mutations (or the area under the intensity curve), is a trade-off between these opposing treatment effects.

**Fig 2.**
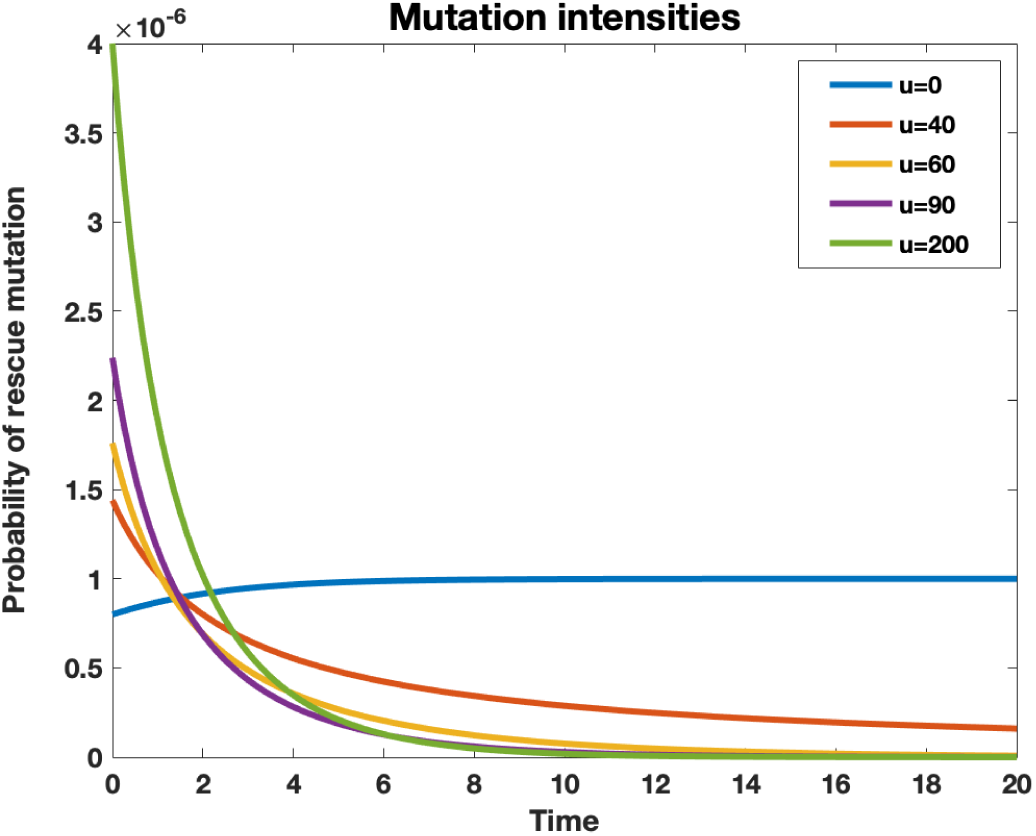
Mutation intensity profiles. Each treatment strategy *u*(*t*) leads to a characteristic mutation intensity profile *S*(*t, u*(*t*))*µ*(*u*(*t*)), which gives the probability of gaining a rescue mutant as a function of time. If no treatment is administered (*u*(*t*) = 0), the population grows to its carrying capacity and generates rescue mutations at a constant baseline mutation rate (blue). A dose-dependent mutation rate introduces a trade-off, where treatment can be used to decrease the population size only at the expense of simultaneously increasing the mutation rate. This creates a sharp mutation peak at the beginning, when treatment is applied to a large population size. The optimal treatment strategy, which minimises the probability of an evolutionary rescue, exploits this trade-off by balancing the early mutational peak such that the area under intensity profile is minimised.

For evolutionary rescue to occur, the time at which the rescue mutation emerges plays no role. Early and late mutations are considered equally bad if the clinical objective is to maximise the probability of a complete cure. However, if evolutionary rescue does occur then the time of mutation is integral in determining the expected rescue fraction. This is simply because resistant cells that emerge early during the treatment period can generate much more growth than resistant cells that occur late. To minimise the expected number of resistant cells at the end of the treatment period, we need to weigh each mutation by the growth it can generate. Assuming a simple exponential growth of the resistant cell population at rate *r*_*R*_, the cost functional needs to be modified with a discount term 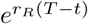, which equals to the growth generated by a resistant cell that emerged at time *t*. We refer to the problem

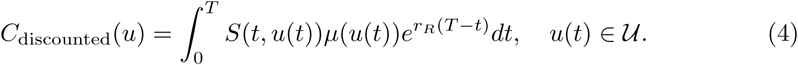

as the discounted control problem and show that the strategy that minimises the expected number of resistant cells at the end uses even lower doses to further reduce the early mutational peak.

### Intermediate dosages become optimal already with modest dose-dependency

When the perturbed growth dynamics and the dose-dependent mutation rate are specified, the optimal treatment strategy, which minimizes the chosen objective, can be calculated using the methods of optimal control theory. As the cost functional (3) depends only on the sensitive cells (and there are initially so few resistant cells present that their competitive effect on the sensitive cells is negligible), we can use deterministic dynamics to calculate the mutation intensities. Using a logistic growth model with Hill-type pharmacodynamics and linear dose-dependent mutation rate (Eq. 7, Materials and Methods) we first solved the optimal control problem (3) using the Forward-Backward Sweep Method [27], which is based on Pontryagin’s minimum principle. The optimal control strategy *u*(*t*) together with the optimally controlled trajectories are shown in Fig. 7. Resistance will always emerge when using the deterministic dynamics, because partial mutations are generated during each time step. We further performed stochastic population dynamics simulations to gain a more realistic depiction of resistance evolution, as discussed later.

**Fig 3.**
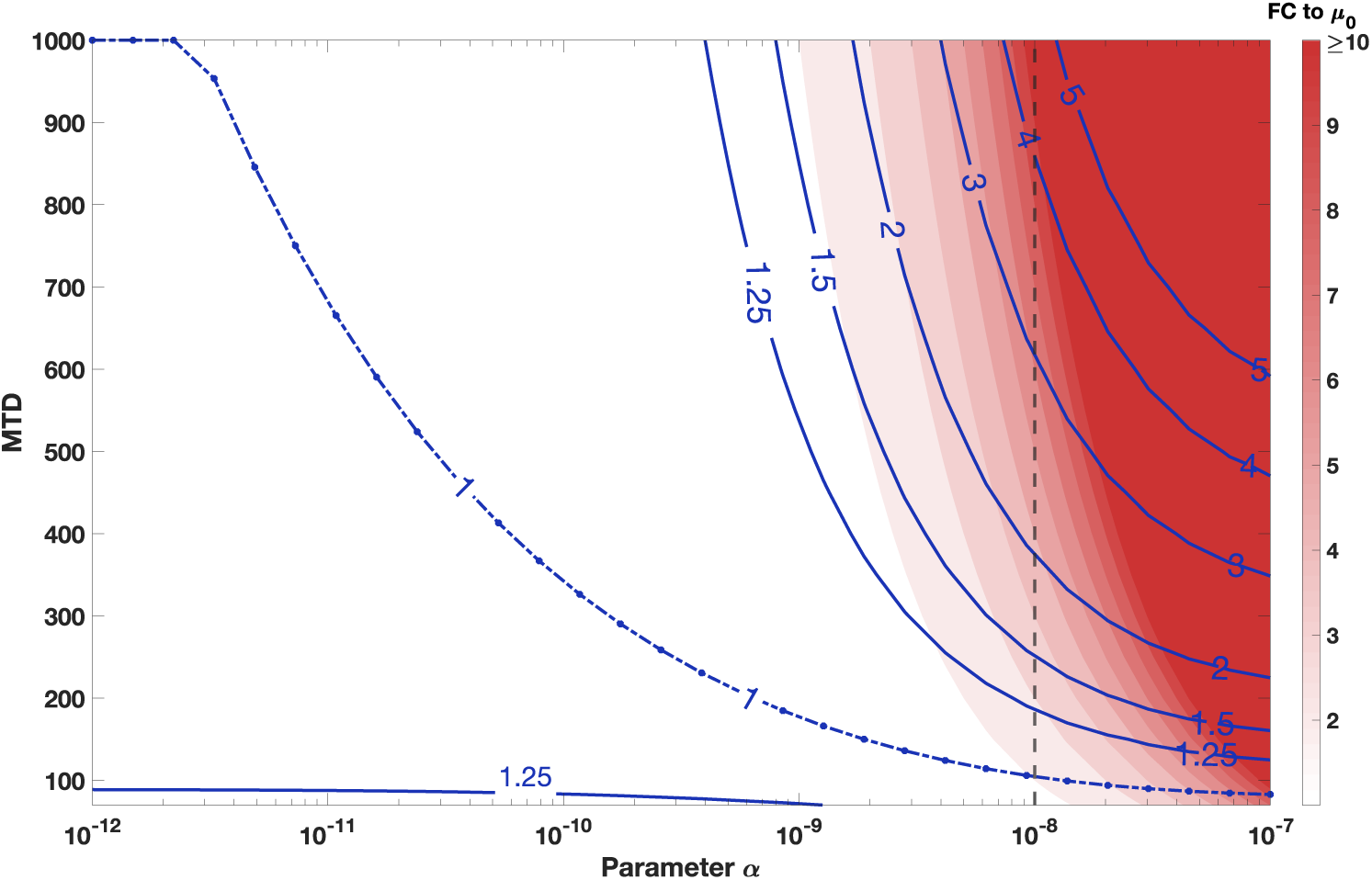
Optimal therapies substantially reduce the number of resistance mutations generated compared to MTD. The blue contour levels correspond to the cumulative mutation intensities (the expected number of rescue mutants) relative to the optimal constant treatment (the dashed 1-line). For example, the 2-line gives the cases where the corresponding MTD produces 100 % more rescue mutations than the optimal dose. The red background color indicates the assumed fold change (FC) to the baseline mutation rate. We notice that substantial improvements are possible even for modest fold-changes depending on how well the drug is tolerated. The probability of evolutionary rescue scales exponentially in the amount of the rescue mutations. We consider the case *α* = 10^*−*8^ in detail, which corresponds to the cases given by the vertical dashed line.

**Fig 4.**
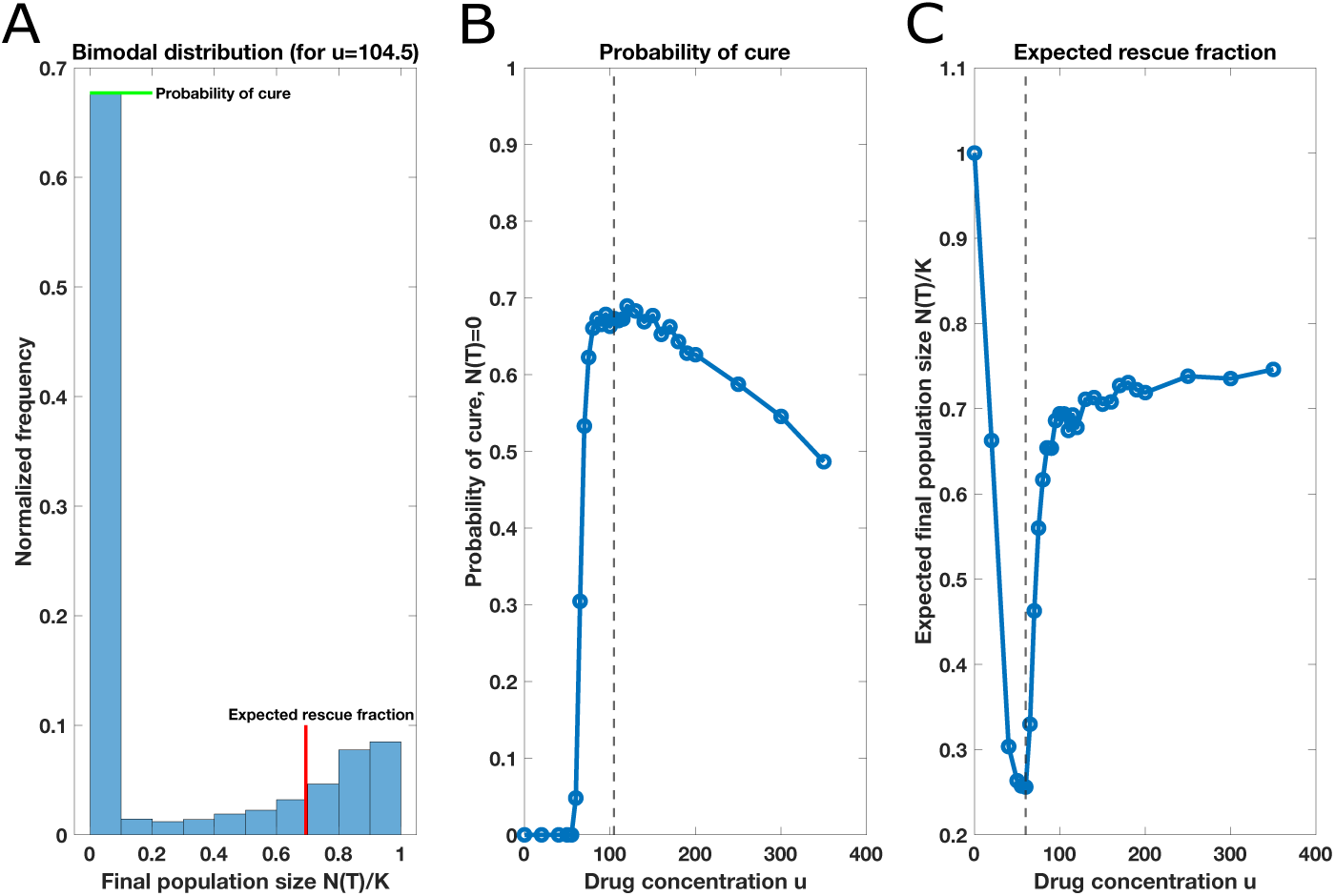
Non-monotonic dose responses. *N*_sim_ = 2000 constant therapies were simulated for each dose while recording the final population sizes. **A** Example of a bimodal distribution of the final population sizes using the optimal constant dose *u* = 104.5 that minimises the rescue probability (cost function Eq. 3). The zero mode corresponds to the proportion of extinct populations (cure) and the second mode corresponds to the expected rescue fraction conditioned on non-extinction. Each dose leads to its own characteristic bimodal distribution. **B** The proportion of extinct populations *N* (*T*) = 0 plotted as a function of dose. The probability of cure displays non-monotonicity and is maximised in the neighborhood of the optimal dose *u* = 104.5 (dashed line) as expected corresponding to cost function given in Eq. 3. **C** The expected rescue fractions *N* (*T*)*/K* conditioned on non-extinction (extinct populations were excluded) plotted as a function of dose. The expected rescue fraction is minimised at *u* = 60 (dashed line), which is the minimising solution of the discounted cost function Eq. 4.

**Fig 5.**
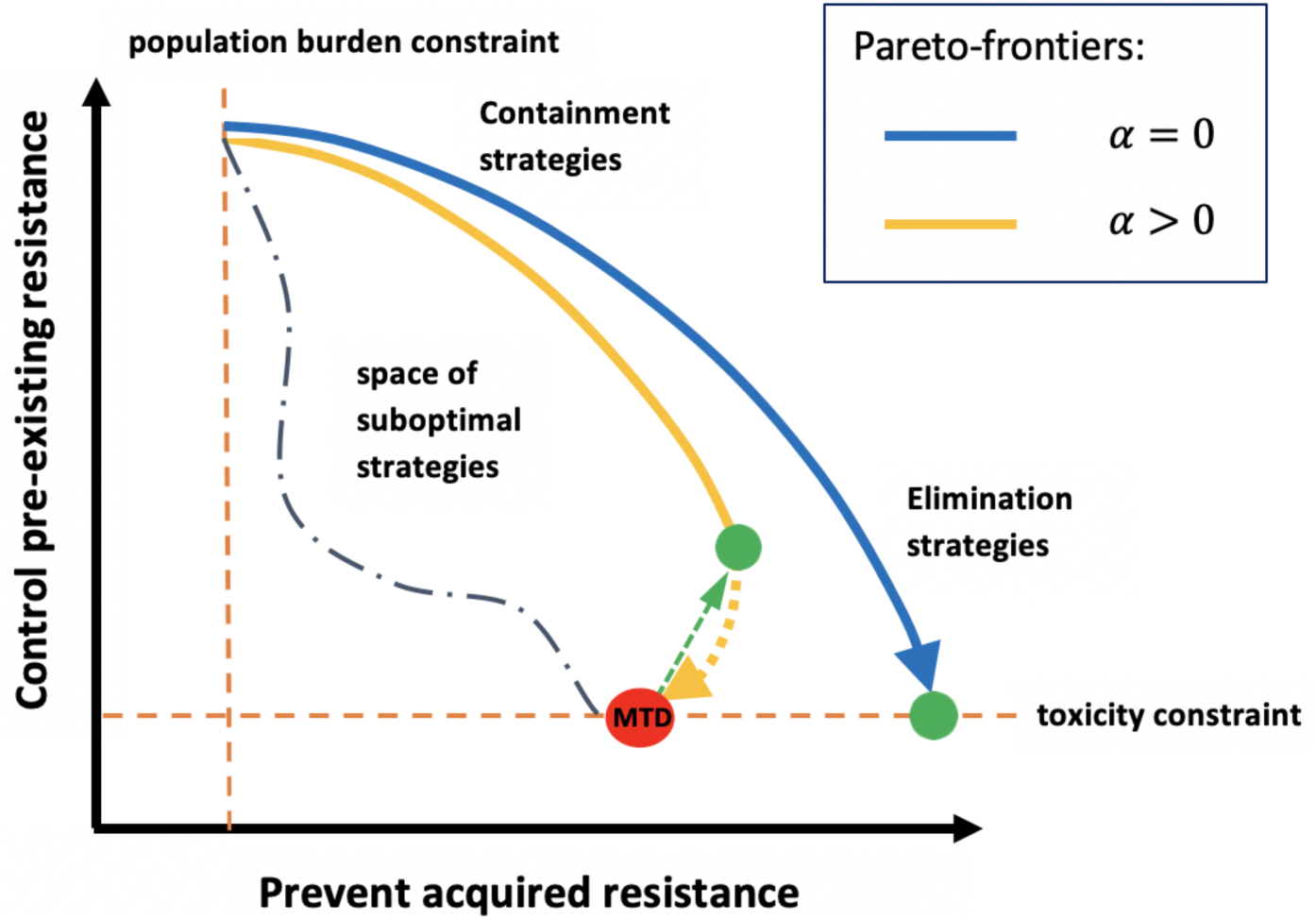
Trade-offs in treatment optimisation. Every treatment strategy is necessarily a trade-off between preventing acquired resistance, by decreasing the population size, and suppressing pre-existing resistance, by allowing intercellular competition. The rate at which the population size can be decreased is constrained from above by the toxicity constraint as well as by finiteness of control leverage and, on the other hand, from below by the population burden constraint, which forces to apply control to stabilize the population size at some acceptable level. When no drug-induced effects are present (*α* = 0), the optimal treatment strategy is found somewhere on the blue Pareto-frontier; the arrow points to the direction where the cumulative drug concentration increases and the optimal elimination strategy (the green point) is given by the MTD-strategy. However, if drug-induced effects are present (*α >* 0), the optimization must be done on a completely different, yellow Pareto-frontier, which exhibits a bifurcation point after which increasing the cumulative drug concentration becomes detrimental with both respects. In these cases, the optimal elimination strategy (the green point) is reached at intermediate dosages at the bifurcation point, which can be identified using the methods presented here. Hence substantial Pareto-improvements (represented by the green arrow) may be achieved by switching from the MTD-strategy to the optimized treatments.

**Fig 6.**
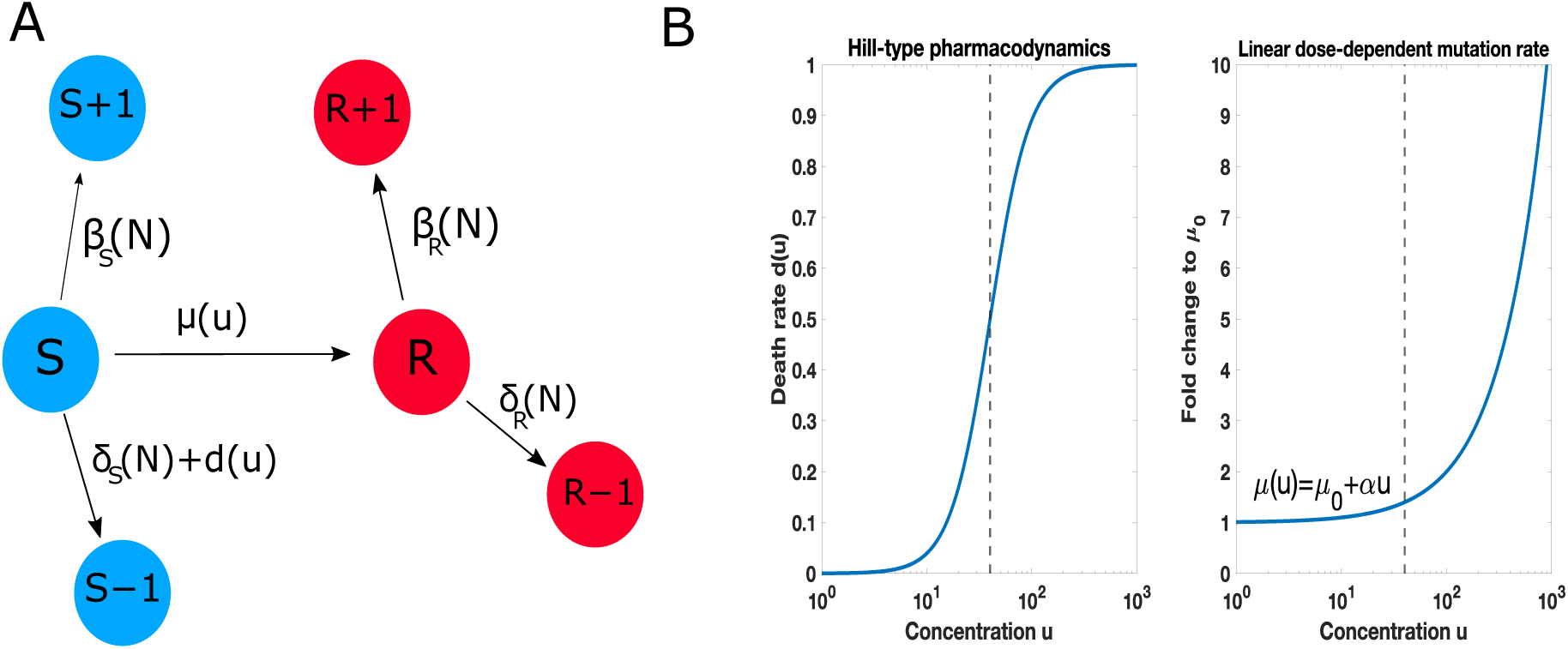
Schematic illustration of the model. **A**. A minimal model for drug resistance distinguishes sensitive (*S*) and resistant (*R*) cells, which follow their own birth-death processes. A treatment can be used to target sensitive cells, but sensitive cells can become resistant via rescue mutations. **B**. Specification of dose-dependent death and mutation rates. We analyse a Hill-type pharmacondynamics and linear dose-dependent mutation rate, but any pharmacodynamics with finite control leverage and monotonically increasing mutation rate will lead to qualitatively similar results reported here. The dashed lines denote the growth inhibitory drug concentration.

**Fig 7.**
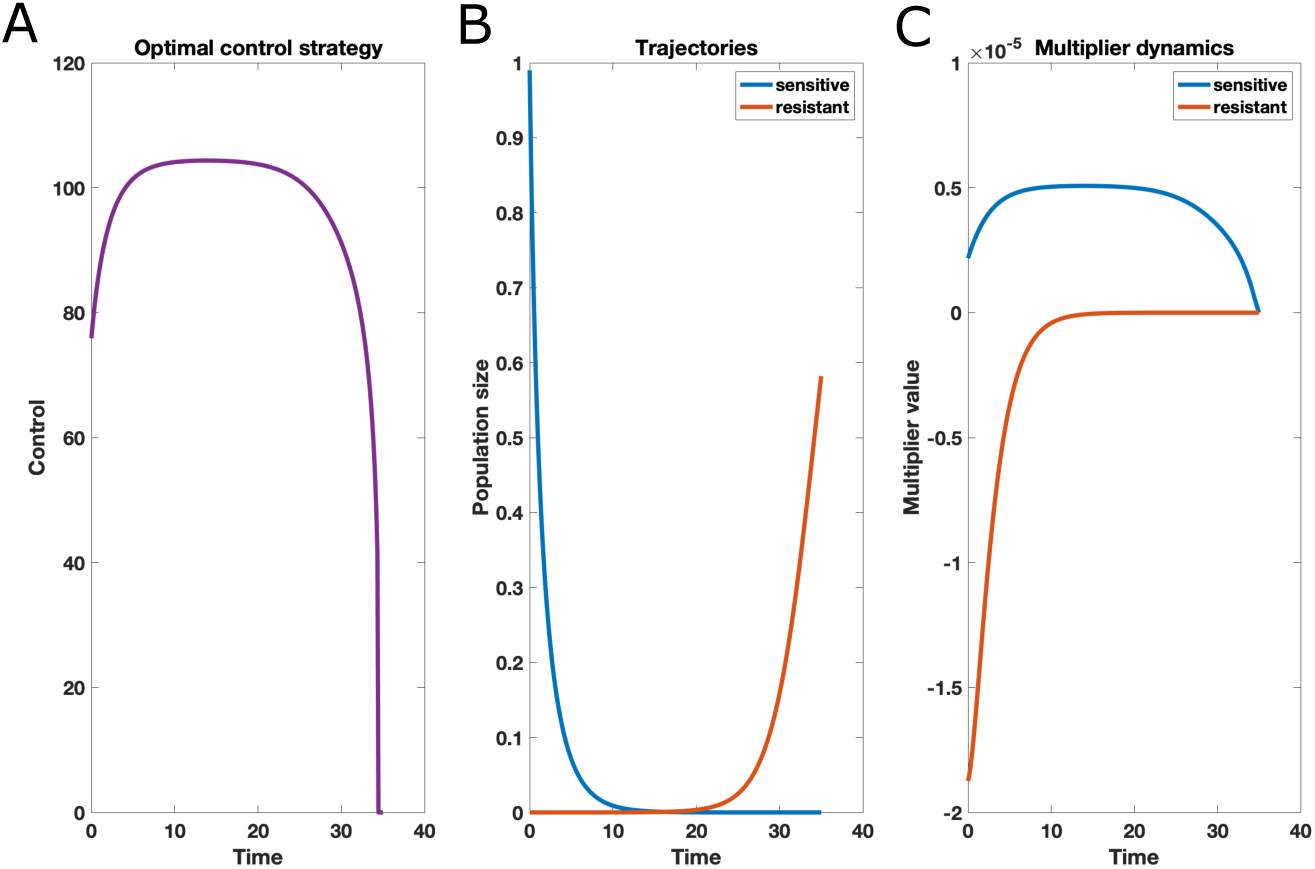
Numerical solution of the optimal control problem of minimising the cost function of eq. (3) using Forward-Backward Sweep Method with parameter values specified in Table 1. **A** Optimal treatment strategy *u*_opt_(*t*) as a function of time. **B** The optimally controlled trajectories *S*(*t*) and *R*(*t*). In deterministic dynamics the population always experiences an evolutionary rescue. **C** The dynamics of the multipliers *λ*_1_(*t*) and *λ*_2_(*t*). The multiplier values can be interpreted as sensitivities of the optimal cost *C*(*u*_opt_) to the perturbations in the respective state variables; here *λ*_2_(*t*) is negative throughout the treatment period, because having more resistant cells would marginally reduce the chosen cost function via competition. Containment strategies become optimal, when the objective function is such that the multiplier corresponding to the sensitive cells becomes negative; it is in these cases that the cost of treatment reduces if there would be more sensitive cells (and therefore more competition) present.

To gain further insights, we solved the same problem using an alternative approach based on the Hamilton-Jacobi-Bellman equation. The resulting control map *u*(*S, t*) (Fig. 8, Supporting Information) is explicitly time-dependent only at the end of the treatment period, which is a boundary effect due to the fixed end time. Therefore, the results are insensitive to the precise time implementation provided the end time *T* is sufficiently large such that sensitive cells can be eliminated during the treatment. In these cases, we can solve for a closed-loop control law *u*(*S*), which depends only on the current population size (Fig. 9). If the treatment period is shorter, the precise implementation time becomes important as the optimal treatment strategy will switch to use no control towards the end.

**Table 1.** Table of parameters used. ([*t*]=unit of time, [*u*]=unit of drug concentration, [*N*]=unit of population density)

| Symbol: | Meaning: | Value: |
| --- | --- | --- |
| $\mu_0$ | baseline mutation rate | $10^{-6}/[t]$ |
| $\alpha$ | slope coefficient of $\mu(u)$ | $10^{-8}/([u] \cdot [t])$ |
| $b_S$ | intrinsic birth rate of sensitive cells | $0.8/[t]$ |
| $d_S$ | intrinsic death rate of sensitive cells | $0.3/[t]$ |
| $r_S$ | intrinsic growth rate ($\beta_S - \delta_S$) of sensitive cells | $0.5/[t]$ |
| $b_R$ | intrinsic birth rate of resistant cells | $0.5/[t]$ |
| $d_R$ | intrinsic death rate of resistant cells | $0.1/[t]$ |
| $r_R$ | intrinsic growth rate ($\beta_R - \delta_R$) of resistant cells | $0.4/[t]$ |
| $d_{\max}$ | maximum death rate by treatment | $1.0/[t]$ |
| $u_{\max}$ | maximum drug concentration | $10^3 \cdot [u]$ |
| $k$ | Hill-coefficient of $d(u)$ | 2.3 |
| $h$ | drug concentration yielding 50 % of $d_{\max}$ | $40 \cdot [u]$ |
| $T$ | fixed end-time of treatment cycle | $35 \cdot [t]$ |
| $K$ | scaled carrying capacity | $1 \cdot [N]$ |
| $S_0$ | initial population density of sensitive cells | $0.99 \cdot [N]$ |
| $R_0$ | initial population density of sensitive cells | $0.00 \cdot [N]$ |
| $\theta_S$ | carrying capacity parameter for sensitive cells | $0.8 \cdot [t]$ |
| $\theta_R$ | carrying capacity parameter for resistant cells | $0.5 \cdot [t]$ |

**Fig 8.**
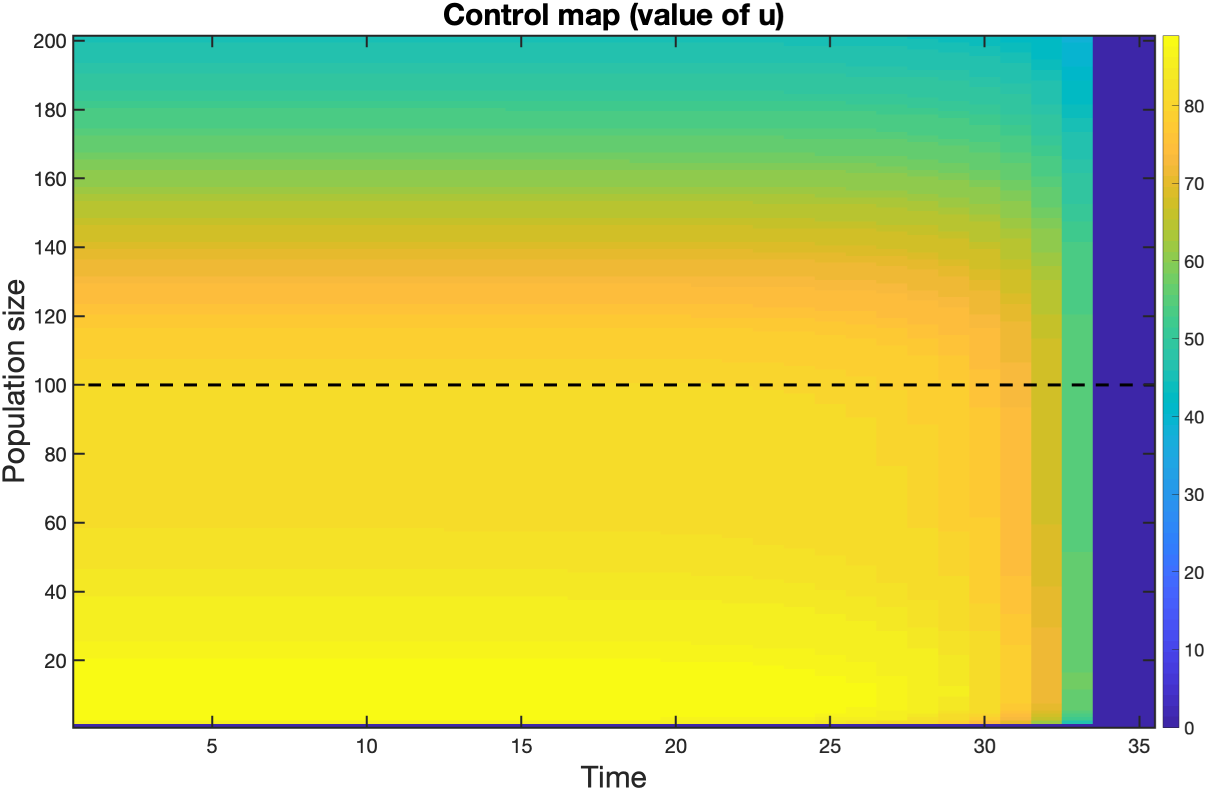
The control map obtained from the stochastic Hamilton-Jacobi-Bellman approach. Here the color denotes the optimal control to be applied at the given population size and time. The carrying capacity has been scaled to *K* = 100. Notice, how the control values start to change in time only at the end of control period, when *t >* 20.

**Fig 9.**
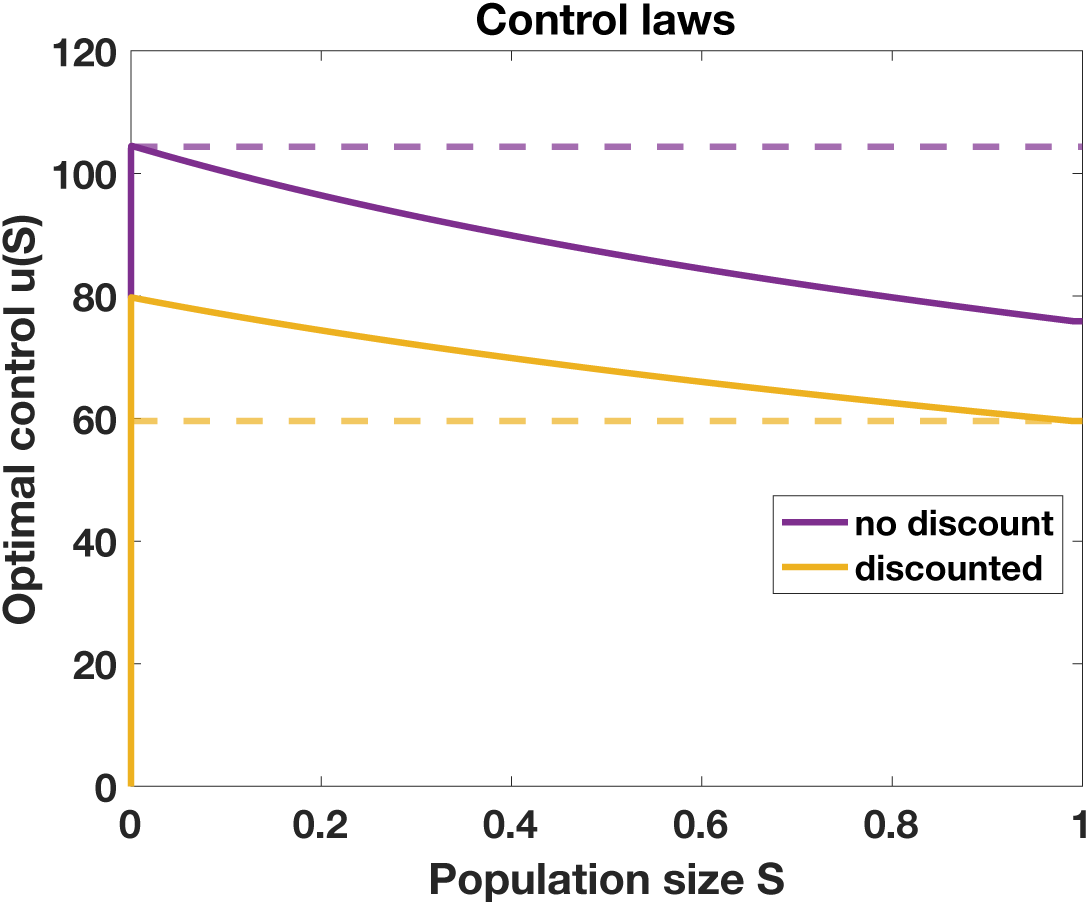
Numerical solutions of the time-independent control laws *u*(*S*) for the problem (3) and the discounted problem (4) using the inverse function method. The dashed lines give the optimal constant doses, respectively. The analytically derived stationary profile matches the numerical solution, but cannot be applied to the discounted problem due to the explicit time-dependence.

The time-independent control law can be derived analytically in an implicit form from the Hamilton-Jacobi-Bellman equation by requiring stationarity (see Supporting Information). As we set the initial resistant cell population to zero and consider only elimination strategies, we can separate the sensitive and resistant dynamics so that the cost functional and the dynamics are independent of the number of resistant cells. For problem (3) with arbitrary density-dependent growth rate *r*(*S*), pharmacodynamics *d*(*u*) and dose-dependent mutation rate *µ*(*u*), we derive the following equation (see Supporting Information)

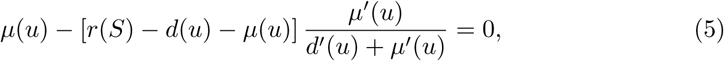

which can readily be solved for the control law *u*(*S*). Notice that here we assumed nothing about the precise functional form of the dose-dependent mutation rate *µ*(*u*), the pharmacodynamics *d*(*u*) or the density-dependent growth model *r*(*S*) except that these are all differentiable functions with respect to *u*. The only technical modeling assumption we have made is that of the *log-kill hypothesis*, where the control leverage depends only of the drug concentration and specifically does not depend on the population size. Therefore, the density dependence realises solely through the assumed density dependent growth rate *r*(*S*), which itself can be approximated to simple exponential decay during treatment. Hence, the optimal therapy is often close to a constant dose and lead us to compare simple constant treatment strategies. Implementing the precise density and time dependencies lead only to marginal improvements and would be more difficult to realize clinically. However, the relative gain of the precise density and time implementation increases, when the drug is less effective (*d*_max_ is smaller) and when the pharmacodynamical profile is less steep (the Hill-coefficient is closer to 1).

Substituting the linear dose-dependent mutation rate and Hill-type pharmacodynamics (see Fig. 6B) in Eq. 5, we obtained the optimal elimination strategy as a function of the parameter *α*, which quantifies the dose-dependency of the mutations, and compared the cumulative mutation intensities relative to MTD. Fig. 3 displays the optimal constant treatment strategy (the blue 1-isocontour) in the relevant region of parameter *α*. Higher doses increase the cumulative mutation intensity or the expected number of rescue mutants generated. As different drugs have different toxicity constraints, there is no universal MTD to compare the optimal dose to. Hence, we regard the MTD as a variable and show few contour lines, where the labels denote the relative cumulative mutation intensity compared to the optimal. The red background color denotes the assumed fold change (FC) to the baseline mutation rate for corresponding *α* and dose.

We notice that the clinical gain of the optimal treatment depends heavily on the drug toxicity; the optimal treatment can lead to substantial gains with already modest drug-induced mutation if the drug is well-tolerated and administered close to, or at, the pharmacodynamical plateau (here *u* = 1000). On the other hand, no substantial gains are achievable even for a highly mutagenic drug if it is also poorly tolerated (i.e., MTD and optimal dosage are close to each other). Here we concentrated our analysis on the case *α* = 10^*−*8^, which leads to only modest fold-change of less than 10 (the vertical dashed line in Figure 3) when compared to the base mutation rate. Following the dashed line reveals that gains on the order of 25 - 100 % are achievable already well below the pharmacodynamical plateau and the maximum dose we considered produces up to almost 5 times more rescue mutations than the optimal. We further note that the rescue probability scales exponentially in the amount of rescue mutations generated (see eq. 2). Therefore, these differences are substantial at the probability level, given that the intrinsic mutation rate is not too large. Indeed, in extreme cases, applying MTD in contrast to the optimal intermediate dosage will switch the emergence of resistance from a rare, mutation-limited, stochastic event to an inevitable outcome.

### Stochastic cell population model demonstrates the efficacy of intermediate dosage therapy under dose-dependent mutation

To further validate our results we performed stochastic simulations and compared various constant therapies in terms of rate of successful eliminations and the size of rescued populations (Fig. 4). The simulations revealed a characteristic bimodal distribution of the final population sizes after treatment, in which the first mode corresponds to extinct populations and the second mode to the expected rescue fraction conditioned on non-extinction. The minimisation of the first mode corresponds to the cost function defined in Eq. (3), where the probability of an evolutionary rescue is minimised, whereas the minimisation of the second mode corresponds to the discounted cost function Eq. (4). Hence, both modes can be analyzed and optimised mathematically using the simpler deterministic dynamics discussed above.

Fig. 4 summarizes the results obtained from stochastic simulation. Fig. 4A displays the bimodality of the distribution of final population sizes. Each treatment strategy leads to its own characteristic bimodal distribution. Fig. 4B shows how the zero mode, that is, the probability of cure, changes as a function of the dose. The different mutation intensities have a substantial impact on the probability of cure and we notice how therapies close to the optimal treatment strategy outperform and lead to substantially better expected treatment outcomes. Similarly Fig. 4C shows an interesting non-monotonic dose response in the expected rescue fraction of target populations that survive from the therapy. The solution of the discounted problem uses even less control in the beginning to shift the expected mutation time to later time points. This of course comes at the expense of increasing the total cumulative mutation rate, and hence, decreases the probability of a cure. This might be acceptable if the probability of evolutionary rescue is in any case high. However, treatment attempting for a cure is always riskier in the sense that if evolutionary rescue does occur, the rescue mutations take likely place very early on thus leading to relapse more quickly. Therefore, it is the baseline expectation of the likelihood of evolutionary rescue which should ideally guide the treatment choice and the precise evolutionary objectives of the treatment.

We conducted the simulations assuming that the emergence of resistance is relatively high but still mutation-limited by setting the effective baseline mutation rate at the start of therapy to *S*(0)*µ*_0_ = 0.1. If the effective baseline mutation rate is even higher, meaning that the emergence of resistance is not mutation-limited, then evolutionary rescue occurs with very high probability in any case, and the relative role of the drug-induced mutations are negligible. However, we would like to emphasize that the viability of *any* elimination strategy relies on mutation limitation, and in these cases, the drug-induced effects become crucially important, and the optimised treatments may lead to substantial improvements as demonstrated in Fig. 4.

## Discussion

MTD therapies are widespread, because the chance of a complete cure is assumed to be maximised by such a regime. This logic dates back to the classical somatic mutation theory of drug resistance, which assumes that resistant cells arise spontaneously at a constant rate irrespective of the treatment. In these cases, the probability of an evolutionary rescue is indeed minimised by eradicating the sensitive cells as fast as possible. Therefore, the MTD paradigm constitutes the optimal treatment strategy when resistance is conferred primarily by *de novo* mutation.

The results obtained herein demonstrate that the situation changes drastically if there are even modest drug-induced effects present. This leads to an interesting trade-off where the population size can be decreased only at the expense of simultaneously increasing the mutation rate. In such cases, the MTD strategy actually increases the likelihood of an evolutionary rescue and thus treatment outcomes may be substantially improved by treatment optimisation. Therefore, it will be of great importance to properly investigate the various mutagenic and other resistance promoting properties of different anti-cancer and antimicrobial therapies in experimental and clinical settings while paying particular attention to dose-dependent mutation rates *in vivo*.

Our main focus was to solve the optimal elimination strategy, which minimises the probability of an evolutionary rescue, in the case of dose-dependent mutation rates. Recently, however, the objective of eliminating the tumor burden has been challenged and so-called containment strategies have been proposed to specifically avoid the competitive release of the resistant cells. Such paradigm shift in treatment may greatly improve treatment outcomes especially in those situations where there is high abundance of pre-existing resistant cells and a complete cure cannot be expected. First proofs of concept have already been made by Gatenby and colleagues in preclinical mouse models and advanced metastatic cancers [32, 33] and recently also in the context of antimicrobial resistance [26]. Based on these findings, Hansen *et al*. argue that all viable treatment strategies must trade-off between minimising mutations (to prevent the emergence of new resistant cells) and maximising competition (to suppress the growth of the existing resistant cells).

To put our results into a wider context, consider Fig. 5 which illustrates this fundamental trade-off between the two alternative evolutionary pressures that can be induced by treatment. So far, the discussion of treatment optimisation has exclusively concentrated on the blue trade-off curve (or Pareto-frontier in the language of multi-objective optimisation), in effect assuming that MTD strategies minimise mutations. The key result obtained here is that this is not generally true when there are drug-induced effects present as the optimisation must then be performed on a completely different trade-off curve. Neglecting the effects of drug-induced resistance can lead to situations where the MTD strategy lies well below the correct trade-off curve and is thus a particularly detrimental strategy as it fails on both aspects. Using the methods presented here, one can identify the optimal mutation-minimising solution and thus potentially gain substantial improvements. Furthermore, the insights gained while studying the discounted problem may be useful also in the context of containment strategies, where partial elimination is sought while lowering the tumor burden to an acceptable level. Thus, when the clinical objective shifts from cure to resistance management, an initial elimination strategy which minimises the expected number of resistant cells becomes a rational objective.

For the case of AMR evolution, our result that intermediate dosage therapy is optimal is particularly interesting as it may also reduce detrimental off-target species effects, such as enriching for resistance in off-target species [12] or compromising community resilience and functioning [34]. Such effects are prevalent in bacterial communities and possibly important to the AMR problem as a whole [12]. As stated above, intermediate dosages may also be optimal in containment strategies [26], which may be useful in chronic infections prevalent owing to factors such as antimicrobial tolerance [35] and biofilms [36] as features of the pathogen population and immunocompromised conditions in the patient. Our findings therefore contribute to an emerging body of evidence showing an increasing scope of utility for intermediate dosages in antimicrobial therapy.

The approach taken here has many advantages. We presented a way of formulating the precise objectives of the treatment in evolutionary terms, which provides an interesting theoretical framework for further treatment optimisation avenues. We specifically considered the effects of drug-induced resistance, an often neglected cost of treatment, and highlighted the importance of these effects. The results obtained provide a qualitative insight that is potentially exploitable by treatment optimisation. Finally, the predicted non-monotonic dose responses of the target populations are also immediately amenable to experimental investigation, where a lot of future study is needed to gain a better quantitative knowledge of the modes and extent of drug-induced effects.

When interpreting the more quantitative predictions made by our work, care must be taken, as they depend on parameter values and more implicit modelling choices (see Materials and Methods for more discussion). For instance, this is the case for the linear dose-dependent mutation rate, where we currently lack extensive data on the dose dependency. This choice was however justified as a plausible and fairly conservative one, motivated by the previous literature (e.g. [14]). Similar qualitative results are likely to be observed for any monotonically increasing dose dependency as nothing in the derivation of Eq. 5 hinges on this particular choice. Also the assumptions related to the growth dynamics and other more implicit assumptions of the dynamical model can be revisited when needed without excessively complicating the solution of optimal controls. Indeed, the purpose of our work was to analyse a minimal model to study the effect of drug-induced mutation instead of providing the most realistic and comprehensive description of the underlying processes. The provided analytical approach can be used to generalise our results to a wide class of models. Given the wide-spread use of MTD therapies, these results may be important and worth of further investigation even if they only apply only to certain drugs.

There remain many outstanding questions for future research related to the topics discussed. Firstly, to be able to design and implement the optimal mutation minimising treatment strategies, we need to have an adequate quantitative understanding of the dose dependency of the mutation rate. Secondly, in the case of containment strategies we need to be able to better characterise and quantify the intercellular competition to which all containment strategies rely on. Thirdly, it is clear that we need to carefully assess and identify those situations where containment strategies would most likely perform better than elimination strategies and how to use both with each other. Finally, identifying and factoring into therapy optimisation the various, as of yet unknown or unquantified, costs of control represent a major goal and unifying principle going forward.

## Materials and Methods

### Dynamical model for drug resistance

Consider the problem of finding the optimal elimination strategy that minimises the probability of evolutionary rescue by *de novo* mutation in the case of drug-induced resistance. First, consider the following general dynamical model for drug resistance:

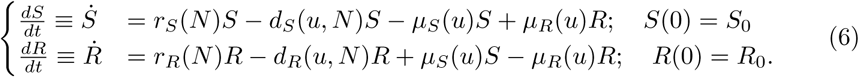

*S*(*t*) and *R*(*t*) are the state variables denoting the population densities for the sensitive and resistant cells, respectively. The functions *r*_*S*_(*N*) and *r*_*R*_(*N*) are the unperturbed growth rates at which sensitive and resistant cells, respectively, grow in the absence of treatment. The growth rate can be different for the sensitive and resistant cells, for example, due to the fitness cost that results from maintaining the resistance mechanism. Constant growth rates lead to the exponential growth model (which is suitable only for small populations) while common density-dependent choices, which depend on the total population size *N* (*t*) := *S*(*t*) + *R*(*t*) via competitive interactions, include logistic 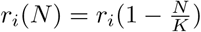 and Gompertz 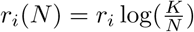 growth models, where *K* is the assumed common carrying capacity and *i ∈ {S, R}*.

The function *d*_*i*_(*u, N*) models the pharmacodynamics of the drug dictating how the obtained concentration of the drug, which is represented by the control variable *u*, translates into cell death. Some drugs can also be cytostatic in nature, meaning that they decrease the birth rate instead of increasing the death rate, which can have important consequences [37] but nevertheless leads to the same mean-field growth as above. Finally, the pharmacodynamical effect may additionally depend on the total population size *N*. By definition, we concentrate on cases where *d*_*S*_(*u, N*) ≫*d*_*R*_(*u, N*)*≥* 0 meaning that resistant cells have a selective advantage during treatment.

Finally, the function *µ*_*S*_(*u*) is the phenotypic mutation rate at which sensitive cells become resistant. Importantly we allow this rate, too, to explicitly depend on the dosage *u*. Furthermore, note that we do not distinguish between the precise cause (genetic and non-genetic) of the change in phenotype, but only consider the transition between the two compartments. Reversible adaptive (epigenetic) changes can be modelled by adjusting the *µ*_*R*_ term.

To demonstrate the qualitative impact of the dose-dependent mutation rate, consider the following simple model with logistic growth, Hill-type pharmacodynamics and linear dose dependency:

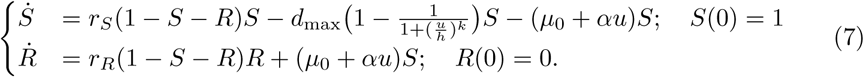

This model follows closely the scaled dynamical model (i.e. *K* = 1) used by Greene et al. but includes more realistic non-linear pharmacodynamics. Now the cost function of eq. (3) can be minimised with respect to the dynamics (7) with two alternative methods based either on Pontryagin’s Minimum Principle (see e.g [25, 38]) or the Hamilton-Jacobi-Bellman (HJB) equation (see e.g. [38, 39]).

Pontryagin’s minimum principle leads to a system of ordinary differential equations that must be solved with mixed boundary conditions, for example, by using the Forward-Backward Sweep Method [27]. The HJB approach on the other hand requires the solution of a partial differential equation for the cost-to-go function, which comes at a higher computational cost. We solve the optimal control problem using both these methods and furthermore provide some analytical insights to the optimal treatment strategy using the stationary profile obtained from the HJB solution.

### Stochastic simulation

To further demonstrate the qualitative impact of the dose-dependency, we performed a stochastic simulation of different constant therapies ranging from low to high concentrations. Each dosage yields its characteristic intensity profile at which rescue mutants are being generated. We show how this connects to the mutational profile and the resulting distribution of rescue fractions that survive treatment.

The stochastic simulation consists of simulating the system given in eq. (7) for a range of constant doses and different initial conditions *S*_0_, corresponding to different effective baseline mutation rates. For each dose, *N*_sim_ = 2000 virtual treatments were administered while recording mutation events and final numbers of sensitive and resistant cells. Unlike the deterministic system, the stochastic birth and death process allows the population to go extinct (a cure). By calculating the proportion of extinct populations for each dose we can estimate the probability of evolutionary rescue and its dose-dependency. Furthermore, by recording the mutational events and stochastic extinctions, we can verify that the stochastic extinction risk is indeed approximately independent of the dose as assumed in cost function of Eq. (3).

For the stochastic system, we need to explicitly specify the birth and death rates and how the carrying capacity is realized (parameter *θ*). The event propensities for the Gillespie algorithm are

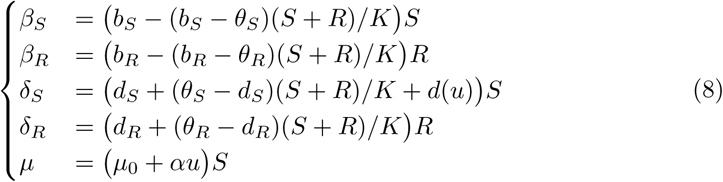

where the events are defined as

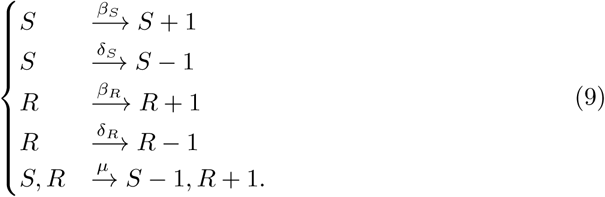

Parameter values that were used in this study are given below in Table 1. We note that the key parameter *α* that sets the drug induced mutation rate slope was selected so that it covers an order of magnitude (see Figure 3 and e.g. [14]). The key elements to observe the discussed trade-off are finiteness of control leverage (with molecular binding based control this is generally true) and monotonically increasing dose to intrinsic mutation rate dependency. The other parameters are chosen such that, the therapy can enforce the sensitive population to decay while the resistant cell can grow once established. These rates then set a sufficient time *T* that ensures that in most cases the sensitive population has been eradicated and the resistant population, if established, occupies a substantial part of the released niche. As discussed earlier the product *S*_0_*µ*_0_ fixes to what degree the evolution of resistance is mutation limited. If that product is large to begin with there is not much help in optimising the therapy induced mutations. In such a case the objective of the treatment should move away from eradication.

### Code availability

The codes used are available from github.com: https://github.com/mustonen-group/drug-induced-mutation/

## Author contributions

**TK** Conceptualization, Formal analysis, Investigation, Methodology, Software, Writing – original draft. **JC** Investigation, Writing – review & editing. **RN** Investigation, Writing – review & editing. **NB** Investigation, Writing – review & editing. **TM** Investigation, Software, Writing – review & editing. **VM** Conceptualization, Formal analysis, Funding acquisition, Investigation, Methodology, Supervision, Writing – original draft.

## Supporting Information

### Solving the optimal controls using Pontryagin’s minimum principle

Consider the optimal control problem

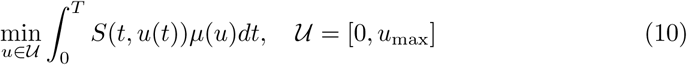

with *µ*(*u*) = *µ*_0_ + *αu*(*t*), subject to the dynamics

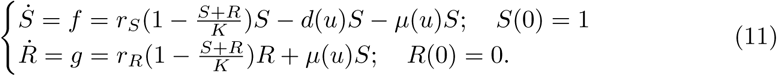

where 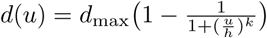 fixing the pharmacodynamics of the drug. The problem can be solved with two alternative methods based either on Pontryagin’s minimum principle or Hamilton-Jacobi-Bellman equation. The Pontryagin’s minimum principle is based on the variational approach and proceeds by defining the Hamiltonian as

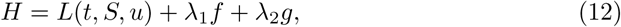

where *L*(*t, S, u*) := *S*(*t, u*(*t*))*µ*(*u*) is the Lagrangian cost functional and the multipliers (costate variables) satisfy the following equations:

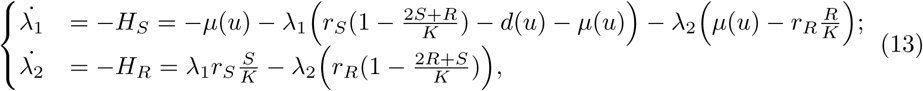

with boundary conditions

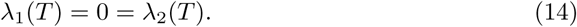

The optimal control strategy *u*_opt_(*t*) is found by studying the function

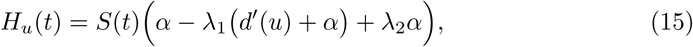

whose roots, should they exist, determine the singular controls. (If no roots exists, then the optimal control reduces to so-called *bang-bang* controls.) Using the parameters given in Table 1, two separate roots appear; the first root can be excluded using the Legendre-Clebsch condition (*H*_*uu*_ *<* 0). The candidate optimal control *u*^*\**^(*S, R, λ*_1_, *λ*_2_) must then be iterated using e.g. the Forward-Backward-Sweep Method [27], where the state variables are first solved *forward* in time using dynamics (11) and then the multipliers are solved *backwards* in time using dynamics (13) with the updated variables.

### Time-independent control laws from Hamilton-Jacobi-Bellman equation

Notice that since the Lagrangian cost functional *L*(*t, S, u*) does not depend on the number of resistant cells *R*, we need not explicitly track them. This is because we assume resistant cells are initially so rare that the competitive interactions on the sensitive cell growth is negligible (note that the opposite is **not** true). If this assumption would be violated, then the cost functional itself would lose its usefulness and the object of the treatment should rather focus in the containment and management of the already present resistance and not its *de novo* emergence. Consequently, we can concentrate our analysis on the single state variable *S* and assume that its dynamics is independent of *R*.

To gain further insight, we then solved the same problem using the alternative method based on Hamilton-Jacobi-Bellman (HJB) equation. This approach relies on solving the *cost-to-go* function *J* (*t, S*) from the partial differential equation:

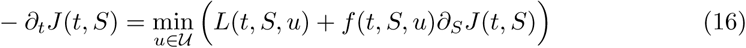

with boundary condition

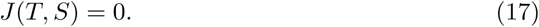

By recording the minimizing control at each point, the HJB approach gives a *control map*

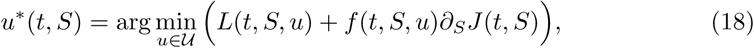

from which the optimal control can be read for any conceivable state. The HJB equation above is for the deterministic version of the control problem that corresponds to the Pontryagin methods described above. In Fig. 8, we solved numerically a stochastic version of the problem.

We notice from Fig. 8 that the optimal control is virtually independent of time except at the very end of the control period, when the control is discontinued. Closer inspection of this reveals that this is in fact a non biologically relevant boundary effect created by the fixed end-time corresponding to stopping therapy and letting the target population grow again. Consequently, by letting the end-time *T* to be sufficiently large so that the sensitive population is almost surely eliminated before the end, then the optimal treatment becomes time-independent and we can solve for a closed-loop control *u*(*S*) which has only feedback from the current population size. Such stationary solution can be obtained analytically for the deterministic HJB by setting

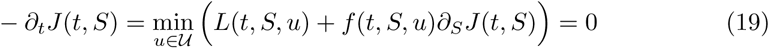

and carrying out the minimisation by formally differentiating the terms inside the brackets with respect to *u*. This minimization yields condition

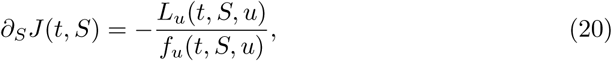

which can be substituted back to the HJB equation. The *stationary profile* for the problem then reads

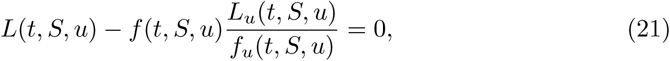

which gives an *implicit* equation for the optimal control. Furthermore, as the time-dependence realizes only via the state and control variables, solving the implicit equation yields a time-independent control law *u*(*S*). Indeed, by substituting the Lagrangian cost functional and the sensitive dynamics, we get

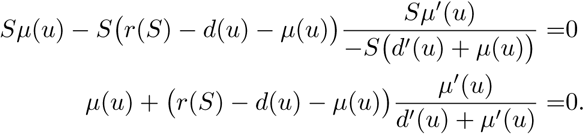

Notice that here we assumed nothing about the precise functional form of the dose-dependent mutation rate *µ*(*u*), the pharmacodynamics *d*(*u*) or the density-dependent growth model *r*(*S*) except that these are all differentiable functions with respect to *u*. Furthermore, notice that as the state variables *S* cancel out, the only remaining dependence of the state variable happens via the assumed density dependent growth model, where we assumed that *r*(*N*)*≈ r*(*S*). Thus, we have now derived equation (5), where the only technical modelling assumption we have made is that of the *log-kill hypothesis*, where the control leverage depends only of the drug concentration and specifically does not depend on the population size.

The analytically derived stationary profile cannot be directly applied to the discounted problem, because of its explicit time-dependence. However, the same stationary profile can be obtained numerically more simply if the optimal time-dependent control eliminates the target population without allowing it to grow in between. Then *S*(*t*) is a monotonically decreasing function and hence there exists an inverse function *S*^*−*1^ : [0, 1]*→* [0, *T*] which gives the time at which the population was at any given size. Now, if indeed time-independent control law *u*(*S*) does exist, it must be unique and thus the optimal control for some population size *S*′ must satisfy *u*(*S*′) = *u*(*t*′) where *S*(*t*′) = *S*′. Then the stationary profile can be obtained using the inverse function as

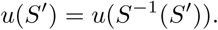

Applying this inverse function method agrees with the analytically derived stationary profile and can be used also to the discounted problem (Fig. 9).

Here we have focused on the problem of determining the optimal way of eliminating the target population, with respect to two biologically meaningful objectives. The derived optimal treatment strategies were calculated with respect to the simple constraint that there is maximum concentration *u*_*max*_ which can be tolerated at each time point; however, the cumulative drug concentration was not constrained. Such constraints can also be easily incorporated to the presented optimal control framework by appending the Hamiltonian with an additional state variable, which enforces the isoperimetric constraint (see e.g. [38]).

